# *Pseudomonas* response regulators produced in an *E. coli* heterologous expression host exhibit host-derived post-translational phosphorylation

**DOI:** 10.1101/2021.12.14.472705

**Authors:** Megan E. Garber, Rodrigo Fregoso, Julie Lake, Anne Kakouridis, Aindrila Mukhopadhyay

## Abstract

In this report, we systematically characterize 32 response regulators (RRs) from a metal tolerant groundwater isolate, *Pseudomonas stutzeri* RCH2 to assess the impact of host-derived post-translational phosphorylation. As observed by distinct shifted bands in a phos-tag gel, 12 of the 24 detected RRs show homogenous mixtures of phosphorylated proteins or heterogenous mixtures of unphosphorylated and phosphorylated proteins. By evaluating the phosphorylation state of CzcR and CopR II under varying assay parameters, we found that changes to pH and exogenous addition of phospho-donors (e.g. acetyl phosphate) have little to no effect on phosphorylation state. By applying protein production conditions that decrease the pool of intracellular acetyl-phosphate in *E. coli*, we found a reduction in the phosphorylated population of CopR II when magnesium was added to the media, but observed no change in phosphorylated population when CopR II is expressed in *E. coli* BL21 (DE3) *Δpta*, a mutant with a metabolic disruption to the acetyl-phosphate pathway. Therefore, the specific mechanism of post-translational phosphorylation of RRs in *E. coli* remains obscure. These findings show the importance of characterizing the phosphorylations state of proteins when heterologously expressed, since their biochemical and physiological properties are dependent on post-translational modification.

## Introduction

Two-component systems (TCS) regulate bacterial responses to the environment. In canonical TCSs, histidine kinases (HK) phosphorylate cognate response regulators (RR), which when phosphorylated perform a function^1^. RRs can be chemically phosphorylated *in vivo* and *in vitro* with small molecule phospho-donors such as acetyl phosphate (AcP), carbamoyl phosphate (CP), or phosphoramidate^2^. Biochemical studies for two-component systems are routinely performed with purified RRs derived from recombinant protein production in *Escherichia coli* BL21 (DE3)^3,4^. These experiments are often supplemented with phospho-donors to ensure RRs are in their phosphorylated state^5,6^.

Of the available phospho-donors, AcP is a metabolic intermediate downstream of glycolysis with an important role in energy generation during periods of starvation^7^. Bacteria, including *E. coli*, can use metabolically generated AcP to phosphorylate natively expressed RRs^8^. Post-translational modification by small molecules is speculated to have physiological roles in TCS regulation^8^. Although post-translational modification during recombinant protein expression is a known phenomenon^9^, the possibility for post-translational phosphorylation of response regulators to occur during recombinant protein production has never been systematically assessed. In this work, we report a systematic characterization of the phosphorylation state of RRs produced in *E. coli* BL21 (DE3) with phos-tag gels^6^.

## Results

We heterologously expressed and purified 32 6XHis-tagged RRs derived from *Pseudomonas stutzeri* RCH2^10^ (GCF_000327065.1), and tested their phosphorylation state by phos-tag gel^6,11,12^ and western blot. Phosphorylated protein loaded into a phos-tag gel will migrate more slowly than its unphosphorylated counterpart (Figure 1A). The phosphorylation patterns for each RR were analyzed by comparing the protein migration within a standard 12% SDS-PAGE gel to the migration within a 12.5% phos-tag SDS-PAGE gel (Figure 1B,C, Supplementary Figures 1,2). Of the 32 RRs we screened, we successfully detected 24, of which 12 exhibited full or partial phosphorylation (Figure 1C). Contradicting our expectation for synonymous phosphorylation patterns between paralogs, which we hypothesized because of amino acid sequence conservation in receiver domains, we observed distinctive phosphorylation patterns in RR paralogs. For example CzcR which is paralogous to CusR, CopRI and CopR II, or QseB I, which is paralogous to QseB II and QseB III, were observed to have unique phosphorylation patterns, implying that the phosphorylation state could be dependent on a physical characteristic that is intrinsic to the RR, despite its homology to other RRs. Overall, initial observations from the high throughput screen demonstrate that the phosphorylation state of a purified RR can be impacted by a recombinant protein expression host.

**Figure 1:**
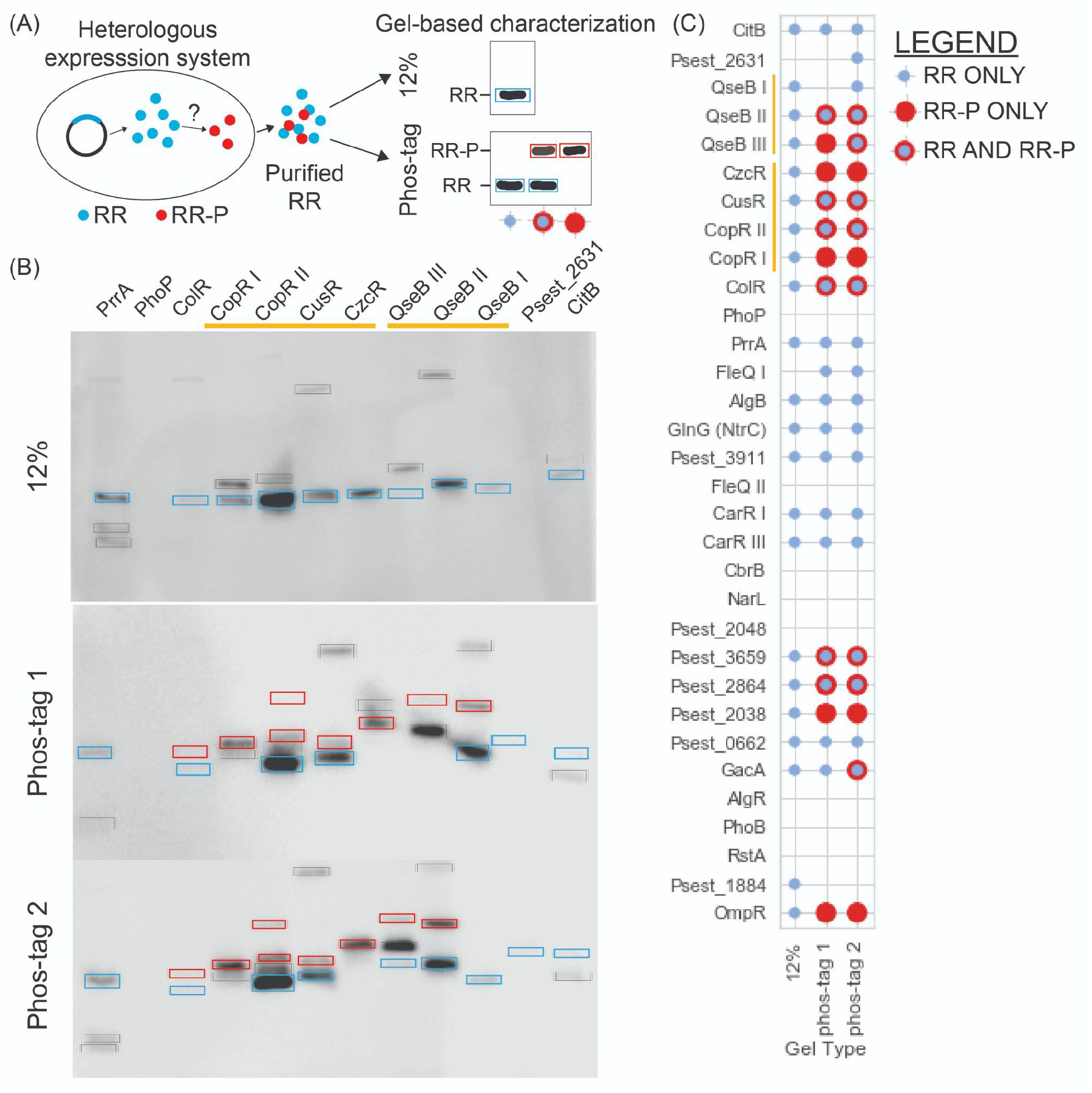
Systematic characterization of phosphorylation state of heterologously expressed Response Regulators (RRs) (A) Methodological overview used to systematically characterize RR phosphorylation state. RR is heterologously expressed in *E. coli*, purified, and characterized on 12% or phos-tag gels. Purified RR will not shift in a 12% gel, however can exhibit three possible patterns in phos-tag gel: unphosphorylated (no shift), partially phosphorylated (1 unshifted band and 1 or more shifted bands), or fully phosphorylated (1 or more shifted bands). (B) 12% and phos-tag gels of select RRs from *Pseudomonas stutzeri* RCH2. Blue boxes around unphosphorylated protein, red boxes around shifted phosphorylate proteins in phos-tag gels, black boxes around ambiguous bands that could be the result of co-purified background proteins or protein dimers. Paralogous RRs referenced in text are indicated by yellow bars. (C) graphical qualification of bands in 12% or phos-tag gels for each RR characterized from *P. stutzeri* RCH2. Blue circles indicate presence of non-phosphorylated bands, red circles indicate presence of shifted, phosphorylated bands, red circles overlaid with blue circles indicate presence of both non-phosphorylated bands and phosphorylated bands. Paralogous RRs referenced in text are indicated by yellow bars.

We re-examined CzcR and CopR II as a case study to understand the contributing factors to the phosphorylation state of RRs during recombinant protein production, and the stability of the phosphorylation after purification. To control for phos-tag gel artifacts, we tested a non-phosphoralatable mutant of CopR II (Figure 2A,B), in which the conserved active aspartate (D51) residue was changed to an inert alanine (D51A). This mutant, which cannot be phosphorylated by cognate HKs or phospho-donors, was found to be fully unphosphorylated when interrogated with phos-tag gel. Importantly, the comparison of the CopR II mutant with its unmutated counterpart confirms that the shifted band observed in the unmutated CopR II is the result of its semi-phosphorylated state. Next, we tested the impact of media composition on phosphorylation patterns. When grown in rich defined media such as autoinduction (AI) media (SF3, ST1) or rich undefined media such as Terrific Broth (TB) (SF4, ST2), both CzcR and CopR II maintain their partially phosphorylated patterns. Then we tested the effect of exogenously adding phospho-donors to the purified RRs (SF3-5, ST1-2). We found that after incubation with phospho-donors the fraction of phosphorylated RR slightly changed, but not dramatically. Next we explored the effect of pH and other dephosphorylation conditions on CopR II (SF3-6, ST1-3). Changes to pH, combinations of treatment with heat, reducing agents, and alkaline phosphatase had little impact on the phosphorylation state. We noted that a fully unphosphorylated population could not be achieved by application of any of these conditions, indicating that the phospho-Asp bond in CzcR and CopR II is relatively stable.

**Figure 2:**
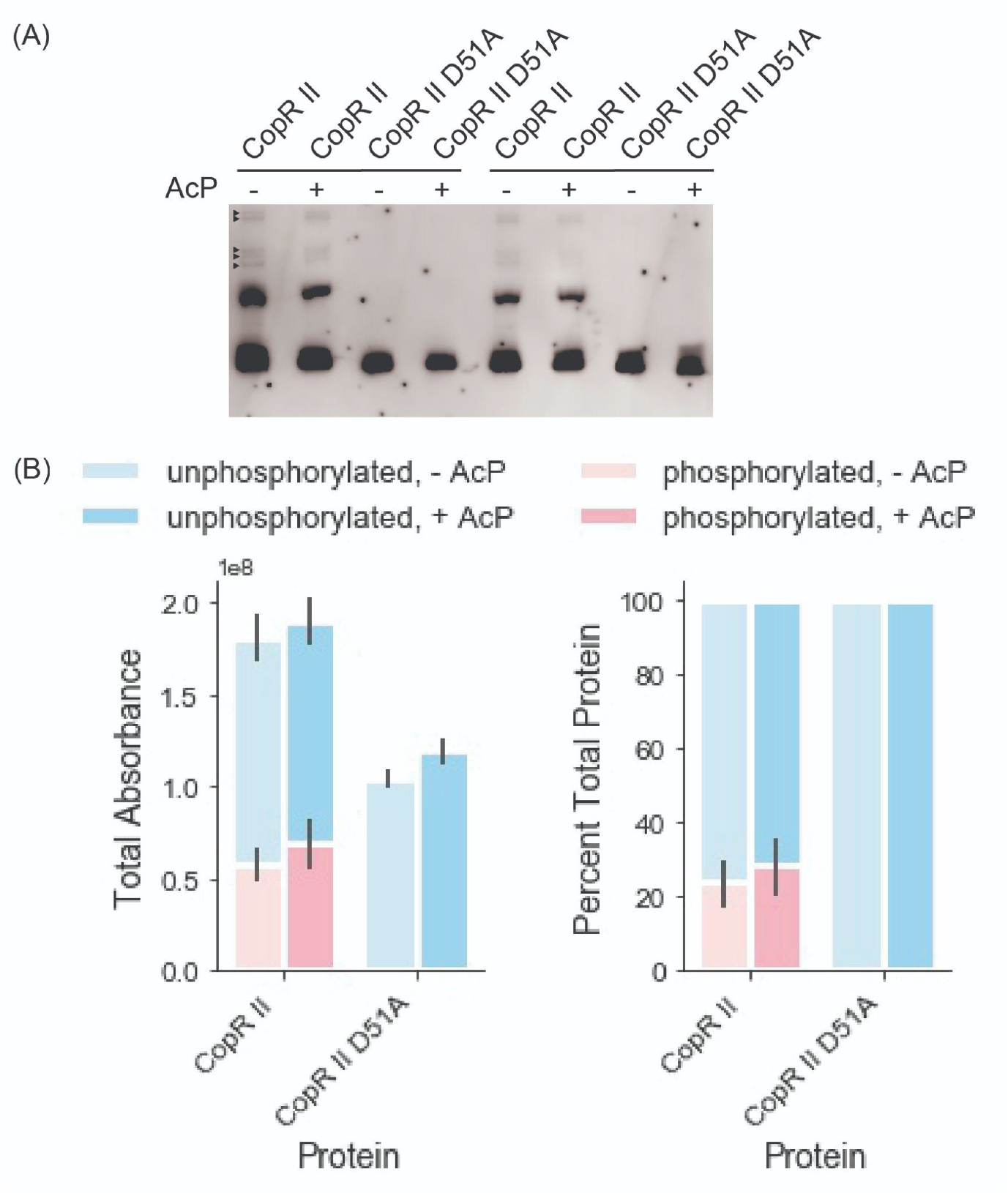
Phos-tag characterization of recombinant CzcR and CopR II. (A) Representative phos-tag result with recombinant CopR II and CopR II D51A. Primary bands were quantified, ambiguous bands that could be the result of co-purified background proteins or protein dimers are marked by small black triangles. (B) Quantitation of unphosphorylated (blue) and phosphorylated (red) bands for CopR II, and CopR II D51A with and without phospho-donor AcP. Bars and error bars represent the averages and standard errors between the average of two technical replicates.

We speculated that the phosphorylation was occurring by post-translational modification within *E. coli*, either by (i) homologs of cognate HKs that could engage in cross-talk with the non-native RRs or (ii) metabolic intermediates that can act as phospho-donors. In *P. stutzeri* RCH2, CopR II is activated in the presence of elevated levels of copper by their cognate HK, CopS^13^. CusS, the homolog for CopS in *E. coli*, is also activated in the presence of elevated copper^14^. To test if *E. coli’s* native CusS was responsible for the observed phosphorylation, we cultivated the CopR II expression strain in the presence and absence of copper (SF7). We found no significant change in the ratio of phosphorylated to unphosphorylated CopR II, indicating that CusS may not play a role in the phosphorylation of the heterologously expressed protein. To test the alternative hypothesis, we cultivated the CopR II expression strain in the presence and absence of magnesium (Figure 3), which has been shown to decrease AcP levels in *E. coli*^*15*^. In agreement with this hypothesis, we observed a decrease in the ratio of phosphorylated to unphosphorylated RR when magnesium was supplemented into the media.

**Figure 3:**
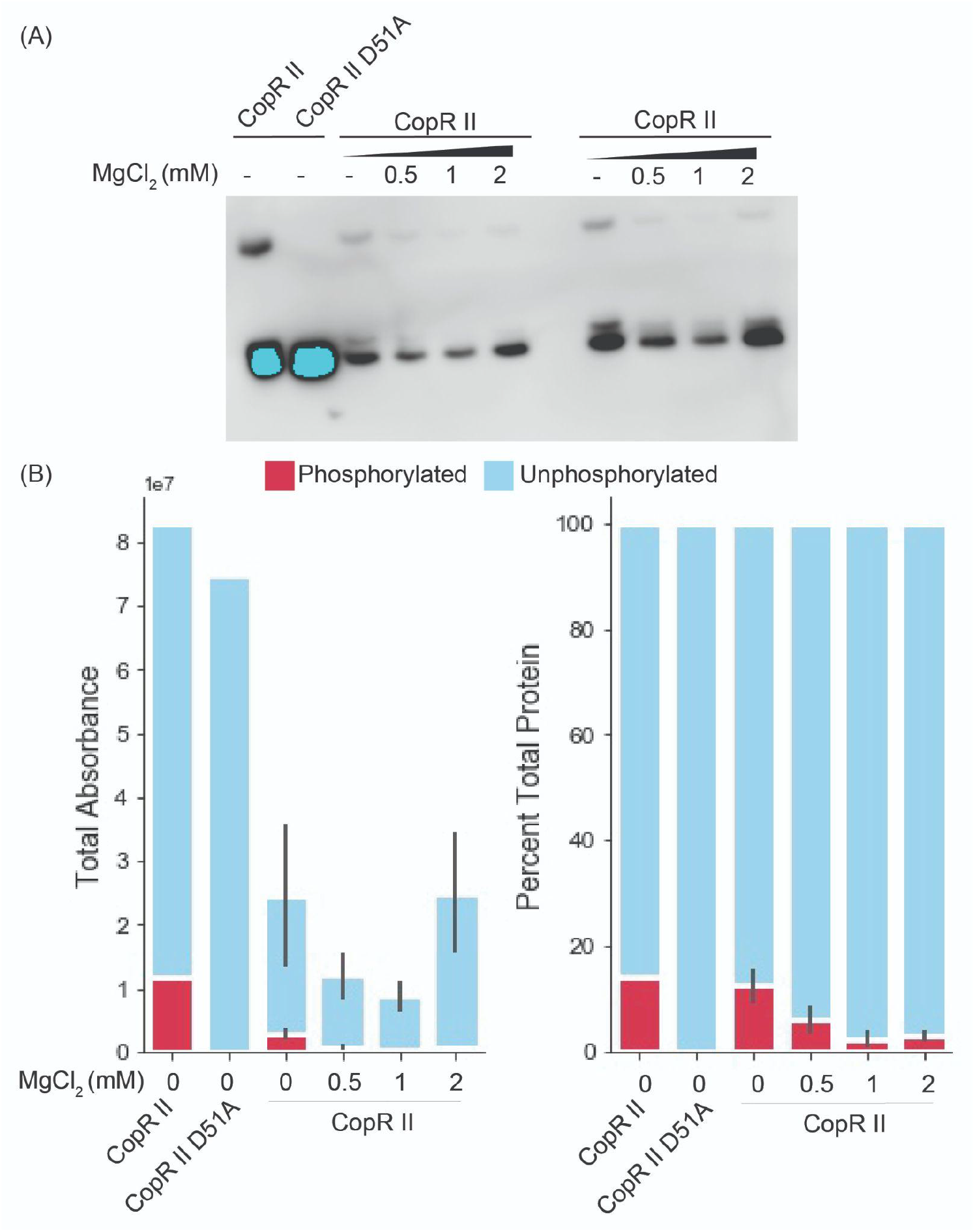
Cultivation with magnesium chloride decreases the phosphorylated population of recombinant CopR II. (A) phos-tag western blot and (B) quantitation by total and percent absorbance for CopR II cultivated in TB media with increased concentrations of MgCl_2_. The first two lanes of CopR II and CopR II D51A were cultivated in a prior batch without MgCl_2_ and were stored at cryogenic temperatures for greater than 2 weeks before electrophoretic separation. Bars and error bars represent the averages and standard errors between the average of two technical replicates.

We next utilized genetics to resolve our hypotheses for the underlying mechanism responsible for the observed phosphorylation patterns. We transformed *E. coli* BL21 (DE3) expression strain mutants, *ΔcusS, ΔackA*, or *Δpta* with the CopR II expression vector and characterized the resultant purified proteins with 12% and phos-tag gels (Figure 4). The amount of phosphorylated RR increased in the absence of CusS, suggesting that CusS acts on heterologous CopR II as a phosphatase. Increasing the AcP levels by deleting *ackA*, marginally increased the phosphorylated population. However, no change was observed in *Δpta*, a strain that has previously been shown to have depleted levels of AcP. Together these results indicate that neither the basal levels of AcP nor the homologous HK are the primary cause of the observed phosphorylation pattern in the parent *E. coli* expression background, however metabolically generated AcP can phosphorylate heterologously expressed RRs.

**Figure 4:**
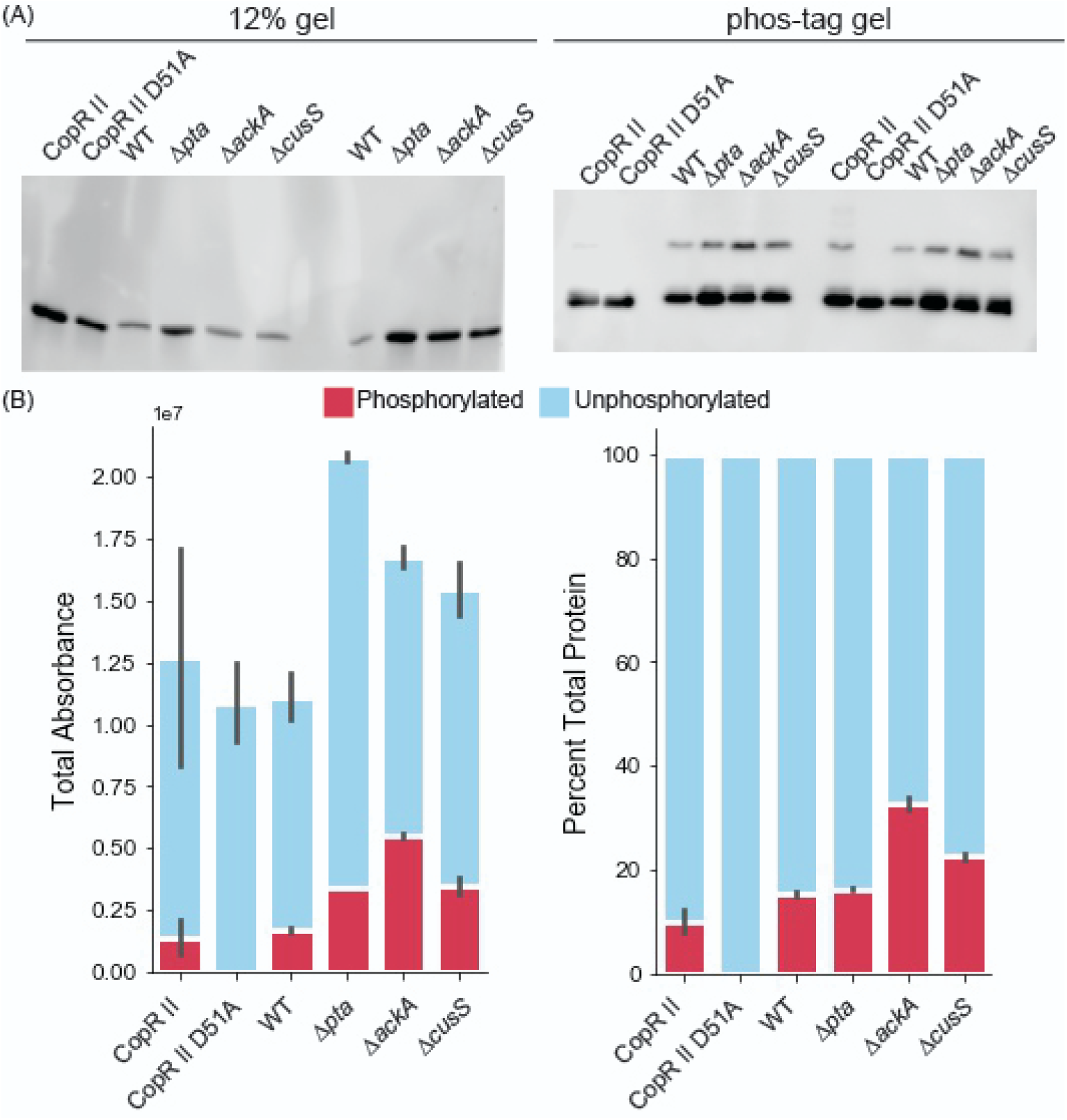
Cultivation in deletion strain influences the phosphorylated population of recombinant CopR II. (A) 12% and phos-tag western blots and (B) quantitation by total and percent absorbance for CopR II cultivated in TB media in *E. coli* BL21 (DE3) (WT) and deletions strains (Δ*pta, ΔackA*, and *ΔcusS)*. The lanes labeled CopR II and CopR II D51A were cultivated in a prior batch in the WT background strain and were stored at cryogenic temperatures for greater than 6 months before electrophoretic separation. Bars and error bars represent the averages and standard errors between the average of two technical replicates.

## Discussion

We demonstrated that RRs can be post-translationally modified in recombinant protein production systems. Because recombinant protein production is routinely used to characterize the biochemical activity of RRs^3,16^, it is possible that the binding and activity rates^17–20^, reported hitherto, could in fact be that of heterogeneous mixtures of RR phosphorylation states. To address this claim, we propose follow-up experiments to examine the binding and activity rates of RRs with homogenous and heterogenous phosphorylation states, determined by phos-tag gel. Because this issue is not yet settled, we suggest that researchers characterize the phosphorylation state of purified RRs of interest by phos-tag gel before biochemical characterization.

It is important to note that phos-tag gels were compared against 12% SDS-PAGE gels to ensure we identified the population of phosphorylated response regulators that only appear in the phos-tag gel, specifically to control for copurified proteins, protein dimers, other ambiguous bands (black boxes in Figure 1B, SF1). Within all of our experiments, the phosphorylation ratios of the proteins remained constant in both technical and biological replicates (Figures 2-4). We note that the phosphorylated to unphosphorylated ratio was quantified using bands with an individual lane, making a standard control accounting for transfer efficiency in the western blots unnecessary. Even though only two technical replicates were performed for an individual experiment, ratios of phosphorylated to unphosphorylated bands remain constant across experiments with different biological replicates (Figures 2-4), showing the reproducibility of our results.

From previous reports on the stability of phospho-Asp bonds in RRs^4,21–25^ we know that despite the short half-life of acyl-phosphate bonds in small molecules^26^, the half-life within RRs varies significantly^1^. In our systematic study, we observed 12 partially or fully phosphorylated RRs. For CopR II, partial phosphorylation was observed even after long-term storage at cryogenic temperatures. Even after treatment with conditions known to destabilize the phospho-Asp bond, we observed that a percentage of the CopR II population remained phosphorylated. In agreement with the previous studies, our study suggests that the lability of the acyl-phosphate bond within RRs is dependent on the structure and conformation of the RR and is not necessarily unstable.

Our study does not reveal the underlying mechanism for the observed phosphorylation of heterologously expressed RRs. First, we do not know how RR structure and homology determine phosphorylation state. Our observation that paralogs of RRs show inconsistent phosphorylation patterns suggests that RR phosphorylation is dependent on intrinsic properties of the RR that are distinct even between paralogs, such as phospho-Asp stability or susceptibility to phosphorylation by small molecule phosph-donors due to surface exposure of the active Aspartate residue. We suggest future studies to determine if orthologous RRs also exhibit distinct phosphorylation trends and structural comparisons of paralogous RRs to evaluate if differences in surface exposure of the active aspartate residue between paralogs is responsible for the differences in phosphorylation. Next, we could not address the mechanism by which RRs are getting phosphorylated. Although our results suggest some interplay between the metabolic intermediate AcP and the recombinant proteins, we could not attribute the observed phosphorylation pattern to its presence. We also cannot rule out the role of homologous cognate HKs in the observed phosphorylation for all of the RRs we tested. However our results with CopR II suggest that we can expect homologous cognate HKs to act as phosphatases when expression strains are grown in non-activating conditions. We propose an alternative hypothesis in which another metabolic intermediate that can serve as a phospho-donor, such as carbamoyl-phosphate, could be responsible for the non-canonical phosphorylation of RRs that are expressed and purified with recombinant protein production systems.

Overall our study provides the phosphorylation landscape for the majority of response regulators for *P. stutzeri* RCH2 as a representative set of RRs that were produced and purified in *E. coli*. Our observations of host-derived phosphorylation demonstrates how valuable it is to test the phosphorylation state of recombinant proteins, however, future studies must examine whether this step is necessary for interpreting subsequent biochemical assay results.

## Methods

### Expression Strain generation

All RRs were cloned into pet28 vectors encoding N-terminal 6X-His tags under T7 inducible promoters. The RRs were PCR amplified from *Pseudomonas stutzeri* RCH2 gDNA with regions homologous to the pet28 plasmid^27^. Parts were assembled via Gibson assembly ^28^ and transformed into *E. coli* DH10b for plasmid propagation. The presence of the insert was confirmed by Sanger sequencing. Plasmids were purified and transformed into expression host *E. coli* BL21 (DE3) or deletion strains if specified. All plasmids generated are listed in ST4.

### Knockout strain generation

A single gene knockout strain of cusS was made in *E*.*coli* BL21 (DE3) according to the Red recombinase method^29^. The gene integration template was prepared using primers previously designed for recombination with cusS (b0570)^30^. Colonies were screened via colony PCR for successful recombination of *E*.*coli* BL21 DE3 Δ*cusS::frt Kan. E*.*coli* BL21 DE3 Δ*ackA:: frt Kan*, Δ*pta::frt Kan* were provided by Alan J. Wolfe lab. All recombination strains were transformed with heat sensitive plasmid, pCP20^30^, at 30 ºC. Single colonies were screened for successful kanamycin flip-out via colony PCR. pCP20^30^ was cured from confirmed knockouts by incubation at 42 ºC. All primers are listed in ST5.

### High-throughput protein purification

Expression strains were grown overnight in LB and subcultured in autoinduction media (Zyp-5052^31^) for growth at 37 °C, 250 RPM, for 5-6 hours and were then transferred to grow at 17 °C, 250 RPM, overnight. Cell pellets were harvested and lysed at 37 °C for 1 hour in a lysis buffer (1X TBS, 100 µM PMSF (Millipore Sigma), 2.5 units/mL Benzonase nuclease (Millipore Sigma), 1 mg/mL Lysozyme (Millipore Sigma). Lysed cells were then clarified by centrifugation at 3214 x g and further filtered in 96-well filter plates by centrifugation at 1800 x g. To enable high-throughput processing, protein purification steps were performed with IMAC resin pipette tips (PhyNexus) using a custom automated platform with the Biomek FX liquid handler (Beckman Coulter). The expressed RRs were individually bound to metal affinity resin embedded within the IMAC resin pipette tips and washed in a wash buffer (1X TBS, 10 mM imidazole). The RRs were then eluted in an elution buffer (1X TBS, 10% (w/v) glycerol, 180 mM imidazole). The purified RRs were then quantified with the QuickStart Bradford Protein Assay (Bio-rad), normalized, flash frozen, and stored at −80 °C for a minimum of one day and up to a week before applying gel-based characterization.

### Low-throughput heterologous expression and purification

Expression strains were grown overnight in LB and subcultured the next day in either autoinduction media (Zyp-5052 ^31^) or Terrific broth (TB) supplemented with Kanamycin (15ug/mL). Cultures grown in TB were induced with 100 µM IPTG at OD600 0.6–0.8 at 37°C and then incubated at 18°C for overnight expression. Cultures grown in autoinduction media were grown for 5-6 hours at 37 °C and then transferred to 17 °C for overnight expression. Cell pellets were harvested and lysed at 37 °C for 1 hour in a lysis buffer (1X TBS, 100 µM PMSF (Millipore Sigma), 2.5 units/mL Benzonase nuclease (Millipore Sigma), 1 mg/mL Lysozyme (Millipore Sigma, Burlington MA)). His-tagged RRs were purified from clarified lysate using HisPur Ni-NTA resin (Thermo Scientific) or HisPur Cobalt resin (Thermo Scientific). The resin was washed in a wash buffer (1X TBS, 10 mM imidazole) and eluted in an elution buffer (1X TBS, 10% (w/v) glycerol, with 250 mM if Ni-NTA resin was used or 180 mM imidazole if Cobalt resin was used). RRs were quantified with the QuickStart Bradford Protein Assay (Bio-rad), normalized, and stored at −80 °C until for a minimum of one day and up to a week before applying gel-based characterization, unless stated otherwise. Biological replicates are defined here as distinct subcultures for protein production. For reactions supplemented with a phospho-donor, 1uL of 1mM Acetyl phosphate was added to 30ug of protein and allowed to react at 37 °C for 1 hour.

### SDS-PAGE (12% TGX or 12.5% Phos-tag gels)

Purified protein was mixed in 1:1 ratio with 2X laemmli loading dye (Bio-rad) prepared according to the manufacturer’s instructions. 10 uL of the protein loading mixture was loaded into the wells of either 12% mini-PROTEAN TGX precast protein gels (Bio-rad) or 12.5% SuperSep Phos-tag precast gels (Wako Pure Chemical Industries). Band separation was achieved by running the gels in Tris-Glycine SDS buffer at 150V for 1-1.5 hours at room temperature. For gels with no ladder CopR II is used as size reference within all gels; the size of all other proteins are measured relative to CopR II, for expected molecular weights see ST4.

### Western blotting and visualization

SDS-PAGE gels were blotted with Trans-Blot Turbo mini 0.2µm PVDF membranes (Bio-rad) using the Trans-Blot Turbo transfer system (Bio-rad) according to the manufacturer’s instructions. After transfer, the membranes were blocked with 50 mL 5% bovine serum albumin in 1X TBS with 0.1% Tween-20 (TBST) for 1 hour. The membranes were then washed 5 times with 1X TBST. Following washing, the membranes were incubated 1 mg L^-1^ of anti-6X-His tag monoclonal antibody [HIS.H8] with an HRP conjugate (ThermoFisher) suspended in 10 mL 1X TBST for 0.5 hours, and washed 3 times to remove unbound antibody with 1X TBST. To activate the HRP conjugate, membranes were incubated with ECL substrate for western blotting (Bio-rad) according to the manufacturer’s instructions. Chemiluminescence was imaged with the Amersham Imager 600 (GE Healthcare) according to the manufacturer’s instructions. Raw image files can be found in the supplemental information (ST6). Two technical replicates of each experiment were performed for each condition. Proteins expressed and purified at different conditions confirm the reproducibility of the observed phosphorylation ratios.

### Image Analysis

Bands were quantified with ImageStudio Lite software (LI-COR v5.2.5). Unphosphorylated and background bands were identified by comparison with images of 12% gels. Phosphorylated and unphosphorylated bands were quantified with the median background method. Background was calculated by estimation of the area to the right and left of the band with a border width of 3 pixels. For images with CzcR and CopR II, all bands in gels and blots were quantified using BioRad Image Lab (Bio-rad v6.0). The background-subtracted densitometry of the bands was determined using multiple band analysis tools within the software. Images were adjusted (contrast, cropping) for representation in main text and supplemental figures.

## Acknowledgments

*E. coli* BL21 (DE3) *ΔackA* and *E. coli* BL21 (DE3) *Δpta* were generously donated by the Wolfe lab.

## Author Contributions

M.E.G. and R.F.O had equal contribution to this work. They performed the experiments, analyzed the results and wrote the paper. J.L and A.K provided technical support. A.M. provided resources, supervision and support. All authors reviewed and edited the final draft.

